# HUGIn: <u>H</u>i-C <u>U</u>nifying <u>G</u>enomic <u>In</u>terrogator

**DOI:** 10.1101/117531

**Authors:** Joshua S. Martin, Zheng Xu, Alex P. Reiner, Karen L. Mohlke, Patrick Sullivan, Bing Ren, Ming Hu, Yun Li

## Abstract

**Motivation:** High throughput chromatin conformation capture (3C) technologies, such as Hi-C and ChlA-PET, have the potential to elucidate the functional roles of non-coding variants. However, most of published genome-wide unbiased chromatin organization studies have used cultured cell lines, limiting their generalizability.

**Results:** We developed a web browser, HUGIn, to visualize Hi-C data generated from 21 human primary tissues and cell liens. HUGIn enables assessment of chromatin contacts both constitutive across and specific to tissue(s) and/or cell line(s) at any genomic loci, including GWAS SNPs, eQTLs and cis-regulatory elements, facilitating the understanding of both GWAS and eQTLs results and functional genomics data.

**Availability:** HUGIn is available at http://yunliweb.its.unc.edu/HUGIn.

**Supplementary information:**

## Introduction

Elucidating the functional roles of the vast majority (>80-90%) of non-coding variants identified from GWAS, and subsequently finding their target gene(s), are pressing and daunting tasks. Investigators have only recently realized the power and value of high throughput chromatin conformation capture (3C) technologies, such as Hi-C (Lieberman-Aiden *et al.*, 2009), TCC (Kalhor *et al.*, 2011) and ChIA-PET (Fullwood *et al.*, 2009), for potential target gene identification (Pombo and Dillon, 2015; Dekker *et al.*, 2017). Several 3D genome browsers, including WashU Epigenome Browser (Zhou *et al.*, 2013) and Juicebox (Durand *et al.*, 2016), have been developed to visualize 3C-based data. However, the vast majority of genome-wide unbiased chromatin organization studies published to date have used cultured cell lines, limiting their generalizability. Understanding chromatin structure and understanding complex trait genetic architecture are each daunting challenges. Understanding both simultaneously is even more daunting, but of paramount importance to unveil genetic mechanisms underlying complex diseases (Gamazon *et al.*, 2015; Jakobsdottir *et al*., 2009).

## Methods

We developed HUGIn (<u>Hi</u>-C <u>U</u>nifying <u>G</u>enomic <u>In</u>terrogator) to host our recently published compendium of Hi-C data across 14 human primary tissues and 7 cell lines (Schmitt *et al.*, 2016) (http://yunliweb.its.unc.edu/HUGIn/). For each tissue or cell line, HUGIn visualizes the observed and expected read counts between two genomic loci, and the corresponding statistical significance, for long range chromatin interactions, in both heatmap and virtual 4C plots (detailed tutorial in **Supplementary Material Section 1** and **Fig. S1**). Compared to existing 3D genome browsers, HUGIn has five major unique features. First, HUGIn simultaneously visualizes a compendium of Hi-C data, which is continually updated as new data become available (**Supplementary Material Section 2**). Second, it directly enables detection of chromosomal organizations both specific to, and constitutive across, tissue(s) and/or cell line(s). Third, HUGIn can display Hi-C data anchored at genomic locus of any size, suggesting potential target gene(s) for GWAS variants in a tissue or cell line informative manner. This feature is particularly important for evaluating a GWAS locus, which can range from a single nucleotide to a region of linkage disequilibrium extending over tens of thousands of base pairs (Smith et al., 2005). Fourth, HUGIn also hosts gene expression and a rich collection of epigenomic data, including typical and super enhancers, CTCF binding sites, frequently interacting regions (FIREs) (Schmitt *et al.*, 2016), and several core histone modifications (**Supplementary Material Section 1**). Lastly, HUGIn generates a list of most likely target gene(s) for any genomic loci of interest. Such a gene list can be used for gene set enrichment analysis or prioritization for follow-up functional studies.

## Results

Figure 1 is a virtual 4C plot in which we display a long range chromatin interaction between the type 2 diabetes (T2D)- and adiponectin-associated SNP rs6450176 with the gene *FST* (Civelek *et al.*, 2017). Although the SNP is located within the *ARL15* gene, it was shown to be an eQTL for *FST* but not *ARL15* in subcutaneous adipose tissue. In addition, expression level of *FST* but not *ARL15* is correlated with adiponectin levels (Civelek *et al.*, 2017). More details and the virtual 4C plot for the rs6450176-FST interaction are shown in **Fig. S2**. Two more examples, a long range interaction between BMI-associated SNP rs9930506 and the gene *IRX3* (Smemo *et al.,* 2014), and between schizophrenia-associated SNP rs1191551 and the gene *FOXG1* (Won *et al.,* 2016), are shown in **Fig. S3-S4**. We further investigated all intergenic SNPs in the NHGRI catalog of published GWAS for seven diseases including schizophrenia, leukemia, Alzheimer’s disease, autism, depression, type 1 diabetes and type 2 diabetes (**Supplementary Material Section 3**). We found that, across all diseases examined, >85% of HUGIn-annotated genes are not the gene closest to the GWAS SNP (**Table S1**); and that HUGIn-annotated gene lists tend to provide stronger evidence for enrichment with biologically relevant terms or gene sets (**Tables S2**-**S3**).

**Figure 1.**
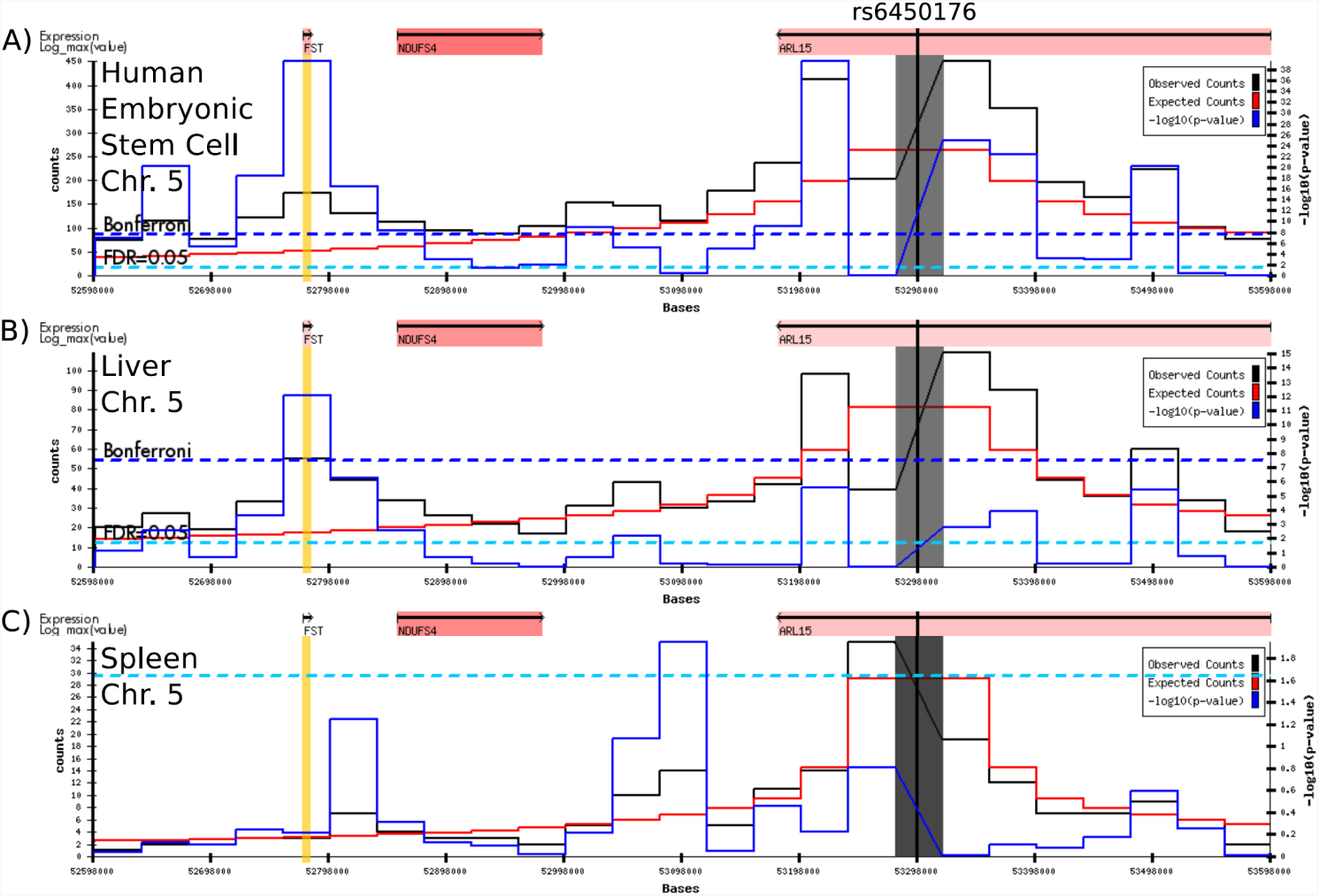
Virtual 4C plot view of the interaction between the T2D- and adiponectin-associated SNP rs6450176 and its potential target gene *FST*. Virtual 4C plot of the long range chromatin interactions anchored at the 40kb bin (chr5:53,280,001-53,320,000), shown as a gray bar, containing the type 2 diabetes (T2D)- and adiponectin-associated GWAS SNP rs6450176 (Civelek *et al.*, 2017), shown as a vertical black line within the gray bar, in H1 human embryonic cell line (**A**), in liver tissue (**B**), and in spleen tissue (**C**). The observed and expected chromatin contact frequency are represented by the black and red lines, respectively. The left Y axis displays the range of chromatin contact frequency. The statistical significance (i.e., — log10(p-value)) of each long range chromatin interaction reported by Fit-Hi-C (Ay *et al.*, 2014) is represented by the blue line, with its range listed in the right Y axis. The cell or tissue specific FDR 5% threshold is shown as a cyan horizontal dashed line, and the more stringent Bonferroni 0.05 threshold is shown as a blue horizontal dashed line. Above each virtual 4C plot is a diagram of genes that are within the 400kb viewing window (chr5: 53,280,001-53,320,000). The highlighted yellow bar is the gene *FST* (chr5: 52,776,264-52,782,304).

## Conclusions

With the continuing accumulation of high resolution 3C-based data in disease-relevant human tissues, we expect that HUGIn will become an increasingly valuable tool for many investigators, including biologists interested in fundamental chromosome organization across cell lines, tissue types, or developmental stages, and geneticists seeking to understand the genetic architecture and mechanism underlying complex diseases and traits.

## Acknowledgements

We would like to thank Dr. Michael L. Boehnke for critical read of an earlier version of this manuscript. We also would like to thank TOPMed investigators for providing helpful comments to improve the usefulness of HUGIn to their research.

## Funding

This work is supported by NIH grants R01HG006292 (YL), R01HL129132 (YL and APR) and U54DK107977 (MH and BR).

*Conflict of Interest: none declared.*

## References

Ay, F. et al. (2014) Statistical confidence estimation for Hi-C data reveals regulatory chromatin contacts. Genome Res, 24, 999–1011.

Civelek, M. et al. (2017) Genetic Regulation of Adipose Gene Expression and Cardio-Metabolic Traits. Am. J. Hum. Genet., 100, 428–443.

Dekker, J. et al. (2017) The 4D Nucleome Project. bioRxiv, 103499.

Durand, N.C. et al. (2016) Juicebox Provides a Visualization System for Hi-C Contact Maps with Unlimited Zoom. Cell Syst., 3, 99–101.

Fullwood, M.J. et al. (2009) An oestrogen-receptor-alpha-bound human chromatin interactome. Nature, 462, 58–64.

Gamazon, E.R. et al. (2015) A gene-based association method for mapping traits using reference transcriptome data. Nat. Genet., 47, 1091–8.

Jakobsdottir, J. et al. (2009) Interpretation of genetic association studies: markers with replicated highly significant odds ratios may be poor classifiers. PLoS Genet., 5, e1000337.

Kalhor, R. et al. (2011) Genome architectures revealed by tethered chromosome conformation capture and population-based modeling. Nat. Biotechnol., 30, 90–98.

Lieberman-Aiden, E. et al. (2009) Comprehensive mapping of long-range interactions reveals folding principles of the human genome. Science (80-.)., 326, 289–293.

Pombo, A. and Dillon, N. (2015) Three-dimensional genome architecture: players and mechanisms. Nat. Rev. Mol. Cell Biol., 16, 245–257.

Schmitt, A.D. et al. (2016) A Compendium of Chromatin Contact Maps Reveals Spatially Active Regions in the Human Genome. Cell Rep., 17, 2042–2059.

Smemo, S. et al. (2014) Obesity-associated variants within FTO form long-range functional connections with IRX3. Nature, 507, 371–5.

Smith, A. V et al. (2005) Sequence features in regions of weak and strong linkage disequilibrium. Genome Res., 15, 1519–34.

Won, H. et al. (2016) Chromosome conformation elucidates regulatory relationships in developing human brain. Nature, 538, 523–527.

Zhou, X. et al. (2013) Exploring long-range genome interactions using the WashU Epigenome Browser. Nat. Methods, 10, 375–6.

